# Understanding the Energy Landscape of Intrinsically Disordered Protein Ensembles

**DOI:** 10.1101/2024.01.04.574178

**Authors:** Rafael G. Viegas, Ingrid B. S. Martins, Vitor B.P. Leite

**Affiliations:** Federal Institute of Education, Science and Technology of São Paulo (IFSP), Catanduva, SP, 15.808-305, Brazil; Department of Physics, São Paulo State University (UNESP), Institute of Biosciences, Humanities and Exact Sciences, São José do Rio Preto, SP, 15054-000, Brazil

## Abstract

A substantial portion of various organisms’ proteomes comprises intrinsically disordered proteins (IDPs) that lack a defined three-dimensional structure. These IDPs exhibit a diverse array of conformations, displaying remarkable spatio-temporal heterogeneity and exceptional conformational flexibility. Characterizing the structure or structural ensemble of IDPs presents significant conceptual and methodological challenges owing to the absence of a well-defined native structure. While databases such as the Protein Ensemble Database (PED) provide IDP ensembles obtained through a combination of experimental data and molecular modeling, the absence of reaction coordinates poses challenges in comprehensively understanding pertinent aspects of the system. In this study, we leverage the Energy Landscape Visualization Method (*JCTC*, 6482, 2019) to scrutinize four IDP ensembles sourced from PED. ELViM, a methodology that circumvents the need for *a priori* reaction coordinates, aids in analyzing the ensembles. The specific IDP ensembles investigated are as follows: two fragments of Nucleoporin (NUL: 884-993 and NUS: 1313-1390), Yeast Sic 1 N-terminal (1-90), and the N-terminal SH3 domain of Drk (1-59). Utilizing ELViM enables comprehensive validation of ensembles, facilitating the detection of potential inconsistencies in the sampling process. Additionally, it allows for identifying and characterizing the most prevalent conformations within an ensemble. Moreover, ELViM facilitates the comparative analysis of ensembles obtained under diverse conditions, thereby providing a powerful tool for investigating the functional mechanisms of IDPs.

## Introduction

Despite the success of Anfinsen’s ideas in understanding the aspects that lead proteins to their respective native states, ^1^ the idea of a single native state is insufficient to understand the functional mechanisms in many cases.^2,3^ There is great plasticity associated with protein functions, often manifesting through multiple conformational states. Understanding the details of these structural transitions is fundamental to comprehending these mechanisms. In terms of structural variability, a significant fraction of proteins (ranging from 15% to 30%) does not have a stable structure, at least when they are not complexed to ligands or binding partners.^4–6^ Such unstructured proteins or protein regions significantly populate a diverse set of conformations and are known as intrinsically disordered proteins or intrinsically disordered protein regions, generally referred to as IDPs and IDRs, respectively. Despite the lack of a rigid structure, IDPs are involved in many cellular functions, ^7–9^ and their great structural variability enables multiple interaction partners exhibiting a wide range of specificity.^10–13^ However, the lack of a rigid native structure leads to significant conceptual and methodological challenges in characterizing structural ensembles of IDPs and IDRs.

The first obstacle is obtaining reliable IDP ensembles. IDPs exhibit intrinsic conformational plasticity, which can be characterized by a rugged and complex energy landscape with numerous, probably shallow, minima.^14^ These characteristics pose significant computational challenges, requiring the use of refined force fields, water models, and sampling techniques capable of efficiently exploring a vast phase space within computationally feasible times.

The strategies used to develop new force fields or to re-parametrize existing ones include the optimization of dihedral angle parameters, strengthening the protein-water interactions, and the introduction of grid-based energy correction map (CMAP) parameters. ^15,16^ For details concerning state-of-the-art force fields for simulating IDPs, refer to recent reviews.^17–21^

Experimentally, a diversity of biophysical methods can be employed to characterize disordered protein states, including Nuclear Magnetic Resonance (NMR), circular dichroism (CD), small-angle X-ray scattering (SAXS), and Förster resonance energy transfer (FRET).^22^ These methods offer insights into the local, intermediate, and long-range structural organizations of IDPs. However, a fundamental limitation arises as these experimental measurements generally reflect average properties, and, in isolation, they are insufficient to uniquely define the underlying heterogeneous ensemble.^23^

One alternative to obtain accurate and trustworthy conformational ensembles is to integrate experimental data with computational techniques.^24–27^ Nevertheless, with this approach, one may often sample averages over too heterogeneous conformations, providing ambiguous information despite being subject to random and systematic errors.^26^ Extensive IDP ensembles curated and validated with experimental data are available at open access on the Protein Ensemble Database (PED, proteinensemble.org).^28^ One can access hundreds of IDP ensembles; however, there is no method capable of providing a general perspective on these ensembles. These conformational sets occur in a highly multidimensional phase space; as a rule, there is no reference structure to define reaction coordinates. To address these challenges, various approaches have been employed in characterizing and comparing ensembles of Intrinsically Disordered Proteins (IDPs). These methods include discrete path sampling,^14,29^ dimensionality reduction techniques, ^30–32^ and the utilization of diverse metrics for assigning distances between different ensembles.^31–35^

One of the possible framework to address this problem is using the energy landscape theory (ELT), which has provided a successful approach to study biomolecules, and in particular protein folding. Based on statistical mechanics principles, it seeks to represent a structural phase space in terms of order parameters, in which the folding funnel concept emerges.^36,37^ The analysis of complex IDP ensembles can be carried out using an approach inspired in the ELT, the Energy Landscape Visualization Method (ELViM). It is a multidimensional projection tool developed to come up with an intuitive representation of the energy landscape of biomolecular systems. ^38,39^ Based on a robust metric, ELViM generates a two-dimensional depiction, in which molecular conformations are optimally arranged according to their pairwise structural similarity. One advantage of ELViM is that its similarity metric is based only on internal coordinates, not requiring a reference state or structural alignment.

In this work, we provide a proof of concept on how ELViM can be employed to systematically tackle and compare IDP ensembles. Given a set of different ensembles of the same system, ELViM can be used to generate an effective phase space encompassing all the systems in the same framework. Analyzing how each individual ensemble populates this entire space may shed light on the sampling heterogeneity of each ensemble, identifying and characterizing the most prevalent conformations within an ensemble. Moreover, this approach allows a comparative analysis of ensembles obtained under diverse conditions. In this study, ELViM is employed to investigate four sets of ensembles that were selected from the PED database. Specifically, the IDP ensembles investigated in this study comprise two fragments of Nucleoporin (NUL: 884-993 and NUS: 1313-1390), Yeast Sic 1 N-terminal (1-90), and the N-terminal SH3 domain of Drk (1-59). Our results show that ELViM can distinguish different ensembles even if the ensembles fit the same averaged experimental data, such as the radius of gyration.

## Methods

### IDP ensembles

All IDP ensembles analyzed in this study were obtained from the PED database, using integrated experimental and computational approaches. We chose available IDP systems for which ensembles were obtained under different conditions. The IDP ensembles analyzed are:

- NUS (1313-1390),^40^ a fragment of the nuclear pore complex protein Nup153 (UniProtKB P49790), available at PED00150. Ensembles are: NUS1 (PED00150:e001, 9482 confs), NUS2 (PED00149:e002, 9405 confs.), NUS3 (PED00149:e003, 9476 confs.), NUS4 (PED00150:e001, 9255 confs), NUS5 (PED00150:e002, 9248 confs), and NUS6 (PED150:e003, 9277 confs.). For ELViM projections, we used one every two conformations.
- NUL (884-993), ^40^ a fragment of the nuclear pore complex protein Nup153 (UniProtKB P49790), available at PED00144. Ensembles are: NUL1 (PED00149:e001, 8003 confs.), NUL2 (PED00144:e002, 8930 confs.), NUL3 (PED00144:e003, 7293 confs.), NUL4 (PED00145:e001, 5938 confs.), NUL5 (PED00145:e002, 6620 confs), and NUL6 (PED00150, 5528 confs.). For ELViM projections, we also used one every two conformations.
- Three ensembles of Sic1 N-terminal targeting domain (1-90).^41^ These ensembles comprise structures of nonphosphorylated Sic1 generated using SAXS (Small Angle X Ray Scattering) and paramagnetic relaxation enhancements (PRE) data (PED00159); SAX, PRE, and chemical shifts (CS) data (PED00160); and an ensemble for the phosphorylated Sic1 N-terminal (at residues Thr2, Thr5, Thr33, Thr45, Ser69, Ser76, and Ser80), generated using SAXS, and PRE. All data was validated using smFRET, for the three ensembles. Each ensemble is composed of 500 conformations.
- Three ensembles of the N-terminal SH3 domain of Drk protein.^42^ In this case, ensembles consistent with NMR, SAXS and smFRET data were generated by the authors using a new approach called Extended Experimental Inferential Structure Determination (X-EISD) method. Specifically, ensembles are: SH3-1 (PED00156, 100 confs.) an unfolded ensemble was optimized from a random pool generated by TraDES;^43,44^ SH3-2 (PED00157, 100 conf.) in which the conformational pool was generated by EN-SEMBLE;^45^ and SH3-3 (PED00158, with 88 conf.) which is a mixed ensemble ranging from disordered to ordered conformations.

### ELViM

The dissimilarity was calculated considering only C*_α_* coordinates. The similarity between two conformations *k* and *l* are defined as

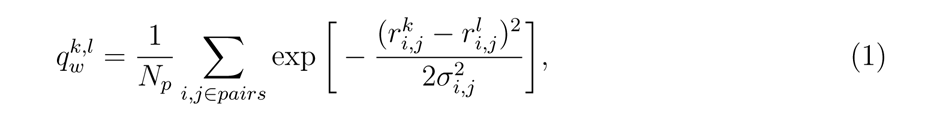

where *r^k^* and *r^l^* are distances between the alpha carbons of residues *i* and *j* in *k* and *l*, respectively. *N_p_* is a normalization constant, and *σ_i,j_* is a weighting parameter that sets the similarity resolution. *σ_i,j_* is defined by^46,47^

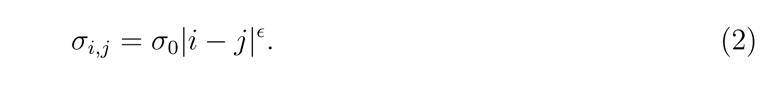

In this study, *σ*_0_ = 3 Å and *ɛ* = 0.15. The dissimilarity between the pair of conformations *k* and *l* is then defined by

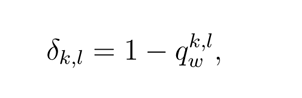

which is unitless, equals zero for two identical conformations, and tends to one for very different conformations.

Considering every pair of conformations within an ensemble results in the dissimilarity matrix used as input in ELViM. The force scheme algorithm^48^ is used to perform a multidimensional projection, which results in a two-dimensional effective phase space. In this representation, each conformation is depicted by a dot, and a heat map is used to map the values of biophysically relevant quantities, such as the radius of gyration, to each conformation. ELViM code is available at github (https://github.com/VLeiteGroup/ELViM). More details about the methodology can be seen in our previous work.^39,49–51^

To characterize the most prevalent conformations within an ensemble, we first estimate the density of data points (density of states) in the ELViM projection. Here, the density was estimated using a Gaussian KDE as implemented in Scipy. ^52^ The KDE bandwidth was adjusted in each case to avoid both over or undersmoothing. High-density regions ideally correspond to the most prevalent conformations within the ensemble and are illustrated using representative conformations, herein called Local Conformational Signatures (LCS). An LCS is found by manually selecting all conformations within a high-density region and finding the centroid conformation, which is defined as the conformation that minimizes the average distance-RMSD. For a detailed description of this procedure, see Ref. ^53^

## Results

To show how ELViM can aid in the interpretation and visualization of IDP ensembles, we selected, as a proof of concept, four different sets of ensembles from the PED. One advantage of the ELViM is that only internal distances are required to calculate the dissimilarities between every pair of conformations. When considering diverse ensembles obtained under varying biophysical conditions or resampled to align with experimental data, ELViM allows for generating a single-effective conformational space, providing a basis for comparing the conformational heterogeneity within each ensemble. Furthermore, this approach facilitates both a differential analysis and a direct visualization of the conformational landscape within each ensemble.

### Structural ensembles of NUS and NUL

The first ensembles to be analyzed are two fragments of the nuclear pore complex protein Nup153 (UniProtKB P49790): NUS (1313-1390), and NUL (884-993). Six ensembles of NUS were analyzed, including three ensembles generated under denatured (NUS1, NUS2, and NUS3), and three ensembles in native conditions (NUS4, NUS5, NUS6). These ensembles were generated by an integrative approach to fit experimental data obtained from SAXs and FRET. More details about the data sets are provided in Ref.^40^ One peculiar characteristic of NUS ensembles is that all six ensembles present approximately the same average radius of gyration (23.38, 23.78, 23.51, 21.45, 21.72, 21.69 Å for NUS1 to NUS6, respectively).

Using ELViM, we can generate an effective phase space, encompassing both native and denatured conditions. For this purpose, we selected one of every two conformations of each NUS ensemble, resulting in a total of 28078 structures. The effective phase space generated by ELViM is depicted in Figure 1. In this representation, structures from all ensembles coexist within the same effective conformational phase space, with each structure visualized as a distinct dot, color-coded based on its radius of gyration (R*_g_*) values. Notably, the radius of gyration values vary smoothly throughout the ELViM projection, with more extended conformations populating the upper border of the projection.

**Figure 1:**
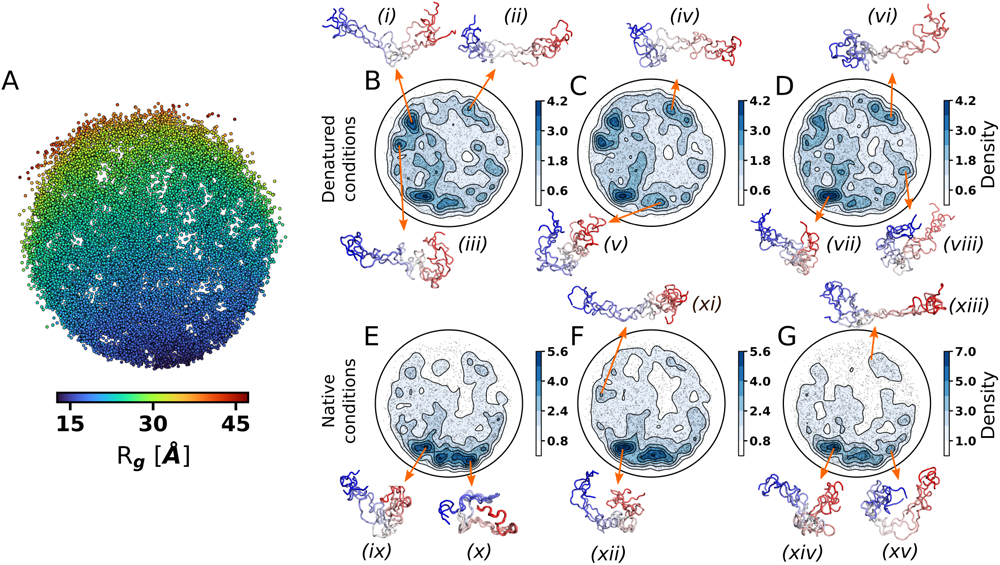
(A) ELViM effective phase space generated for the ensembles of the NUS protein. Ensembles were resampled using one every two conformations, resulting in 28078 conformations. Density of states estimated for the individual ensembles (B) NUS1, (C) NUS2, (D) NUS3, (E) NUS4, (F) NUS5, and (G) NUS6. The backbones of representative conformations from arbitrarily selected regions are displayed with the N-terminus in blue.

Additionally, we are able to separate the conformational phase space for each individual ensemble and compare how they are distributed over the entire phase space. Each configuration of each ensemble can be represented separately in the ELViM projection, as seen in Figure S1. However, due to the superposition of dots in scatter plots, more information can be retrieved by analyzing the density of dots within each ensemble. We calculated the density using a Gaussian kernel (KDE), and the results are illustrated in Figure 1(B-G). This is done for both denatured conditions (B, C, and D in Figure 1) and native conditions (E, F, and G in Figure 1). It is crucial to note that, in unbiased and well-sampled simulations, higher-density regions typically correspond to minima in the free energy landscape; this is not always the case, since bias can enhance the sampling of specific regions of phase space. Nevertheless, ELViM provides an immediate and intuitive visualization of how the sampling is distributed across the entire space for each ensemble. In this case, high-density regions indicate the most prevalent conformations within an ensemble.

Further insights can be obtained by analyzing the conformations within each high-density region. For each individual ensemble, we identify specific Local Conformational Signatures (LCSs) for some of the densest regions as depicted in Figure 1.B–G. In this context, an LCS consists of the centroid structure and its two nearest neighbors, determined by distance-RMSD values. Examining the density of states (Figure 1.B–G), it is possible to note that each ensemble exhibits a unique distribution across the entire ELViM projection. A drastic change in the density profile can be observed when comparing the ensembles corresponding to denatured (Figure 1.B–D) and native (Figure 1.D–F) conditions. Considering only denature or native condition, it is also possible to observe how the density peaks change across different ensembles, showing regions of the phase space that gain or lose stability in the sampling procedure. The similarity in the radius of gyration among these ensembles highlights the utility of such representation, enabling the prompt identification of differences in conformational preferences, even among ensembles that fit the same average experimental observable.

In Figure 1.B–G, we display the density and some LCSs for the individual ensembles of NUS. Under denatured conditions, LCSs *i*, *ii*, and *vii* exemplify regions that are consistently more densely populated across all three ensembles, but with varying density values. Other small basins also gain or lose density in different ensembles, highlighting sampling differences among them. A comparable pattern is evident in the ensembles under native conditions, which populates, with small differences, the lower portion of the ELViM projection. Interestingly, LCS *x* in Fig 1.E corresponds to a region having many well-aligned conformations that loses density for the other two native ensembles, which have more conformational heterogeneity even in the high-density regions.

Each of the densest regions can be further inspected, obtaining the probability distribution contact map and addressing specific structural details. Additional average structural variables of these regions, such as solvent-accessible areas, can be calculated, providing valuable insights into understanding protein mechanisms. ^54^ When one compares the contact maps of similar regions of the ELViM projection for different ensembles, for example regions *ix, xii*, and *xiv* in Figure 1, one can see the most important contacts which are maintained in all native-condition ensembles, Figure S2.

It should be noticed that the ELViM multidimensional projection technique is stochastic, generating slightly different projections for each run (starting with a random distribution of points). Nevertheless, these differences do not impact the global properties of the projection. As an example, we ran two new and independent projection replicas for the NUS system and compared them with the projection shown here (Supporting Information Figure S3). To illustrate the ELViM reproducibility, six groups of points, from different regions of the ELViM projection, were arbitrarily selected in the first replica and shown in the subsequent replicas using the same color scheme. Notably, both the local and global neighborhood are maintained for the three replicas as groups remain cohesive in the same relative orientation (Supporting Information Figure S3).

In a similar approach, three ensembles of NUL were generated under denatured conditions (NUL1, NUL2, and to NUL3), and three ensembles under native conditions (NUL4, NUL5, and NUL6). In the ELViM projection, we selected one of every two conformations of each ensemble, resulting in a total of 21157 structures. Figure 2 illustrates the ELViM projection for the NUL ensembles. Once again, the radius of gyration varies smoothly across the projection. The individual structure representation of each ensemble in the ELViM projection is shown in Figure S4. An intriguing observation is that certain conformations form distinct, well-defined groups, appearing almost detached from the remaining of the projection.

**Figure 2:**
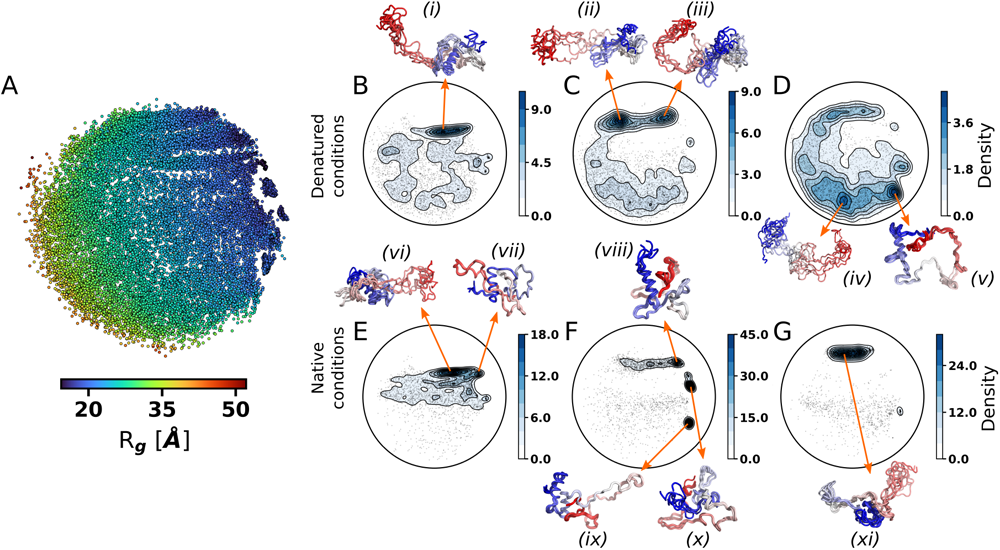
(A) ELViM effective phase space generated for the ensembles of the NUL protein. Ensembles were resampled using one every two conformations, resulting in 21157 conformations. Each dot represents a conformation and is color-coded based on its radius of gyration value. Density of states estimated for the individual ensembles (B) NUL1 (C) NUL2 (D) NUL3 (E) NUL4 (F) NUL5 (G) NUL6. The backbones of representative conformations from arbitrarily selected regions are displayed with the N-terminus in blue.

In Figure 2.B–F, we display the density and some LCSs for the individual ensembles of NUL. When examining the denatured conditions, we display five LCSs labeled from *i* to *v*, each representing distinct dense regions within the conformational phase space. Notably, the densest regions vary among the different ensembles, indicating the diverse conformational behaviors within ensembles generated under denatured conditions. Moreover, although ensembles NUL1 and NUL2 reach their maximum density in the same neighborhood, NUL3 exhibits its maxima displaced to the lower portion of the ELViM projection. Under native conditions, LCSs *vi* to *xi* exhibit significant density, which is confined to smaller regions. Moreover, the three ensembles achieved their density peak in different basins. A reproducibility analysis was also performed for the NUL system and the results are displayed in Figure S5 of the Supporting Information.

### Structural ensembles of yeast Sic1 N-terminal targeting domain

Three ensembles of Sic1 N-terminal targeting domain (1-90) were also analyzed. The ensembles were generated using an integrative modeling and validation procedure.^41^ Namely, ensembles were generated to satisfy different combinations of experimental data from NMR and SAXS experiments and then validated using single-molecule Förster Resonance Energy Transfer (smFRET) data. Specifically: Sic1-1 comprises structures of nonphosphorylated Sic1 generated using SAXS and paramagnetic relaxation enhancements (PRE) data; Sic1-2 is an ensemble generated using SAX, PRE, and chemical shifts (CS) data; and Sic1-3 is the ensemble for phosphorylated Sic1 generated to fit SAXS, and PRE constraints. All three ensembles were validated using smFRET. In this case, we are able to visually compare how different sets of constraint and the phosporilation of some residues can affect the conformational heterogeneity within ensembles.

The ELViM projection generated for these three ensembles, totaling 1500 conformations, is depicted in Figure 3A. In this case, although data points look more sparse, it is possible to observe the formation of local groups (small dense regions) and Rg values also vary smoothly throughout the projection. In Figures 3B, C, and D, projections are colored to show the structures of each individual ensemble for Sic1, Sic2, and Sic3, respectively. The estimated density and some LCSs are also provided for each ensemble. Although the three ensembles seem to be spread throughout the entire ELViM projection, the density analysis reveals details about the most prevalent regions of each ensemble.

**Figure 3:**
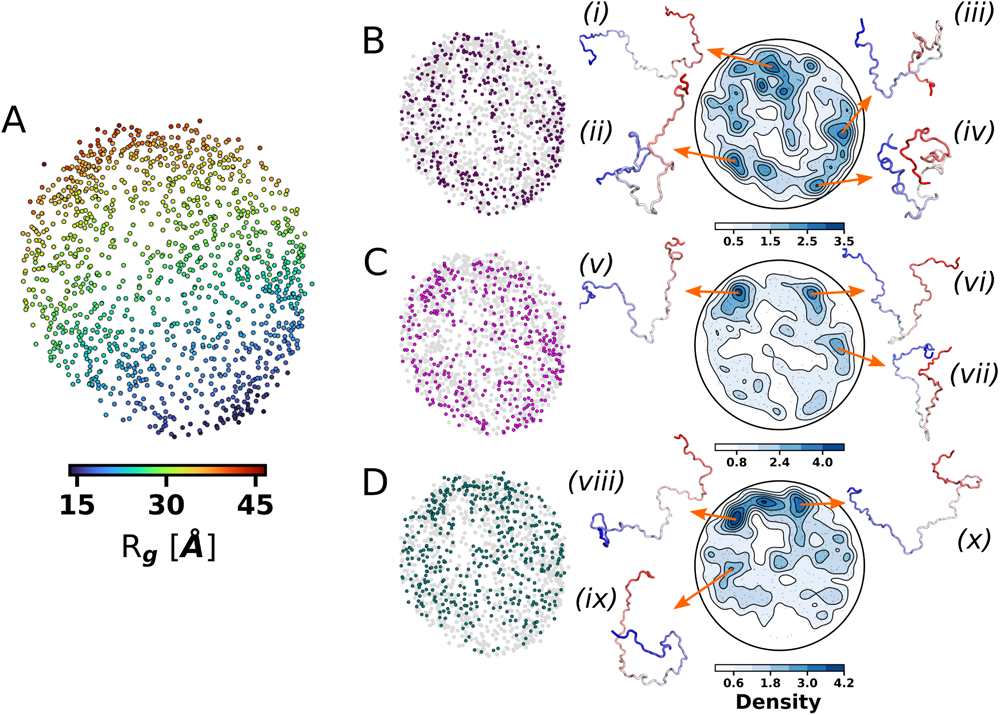
ELViM analysis for the disordered N-terminal of Sic1 (1-90). (A) ELViM projection generated for the three ensembles of Sic1, totaling 1500 conformations. Data points are colorcoded based on Rg values. The conformations of each ensemble are displayed in the middle column in (B) purple, for Sic1-1, (C) fuchsia, for Sic1-2, and (D) teal, for Sic1-3. For each case, a density estimate is also provided, along with representative structures from some high-density regions.

For Sic1-1, which contains structures of the nonphosphorylated protein generated to fit SAXS and PRE data, the high-density basins are more spread throughout the projection than the other cases. The LCSs i, ii, iii, and iv, in Figure 3B illustrate representative conformations with different degree of compactness. Interestingly, for the Sic-2 ensemble (Figure 3C), which was generated to also fit CS data, there are three denser regions on the projections with LCSs that present a similar v-shaped bent in the middle of the structure. In Figure 3D, the same analysis is performed for the ensemble Sic1-3, which is formed by structures of the phosphorylated protein obtained to fit SAXS and PRE constraints. In this case, the denser regions are located at the top side of the projection, where structures of higher radius of gyration are located. The reproducibility analysis performed for the Sic1 ensembles is provided in Supporting Information Figure S6.

### Structural ensemble of the unfolded state of the N-terminal SH3 domain of Drk

Lastly, we analyze three ensembles of the N-terminal SH3 domain of Drk protein detailed in Ref.^42^ In the latter study, ensembles consistent with NMR, SAXS, and smFRET data were generated using the Extended Experimental Inferential Structure Determination (X-EISD) method.^42^ In SH3-1, the unfolded ensemble was optimized from a random pool generated by TraDES,^43,44^ while in SH3-2, the conformational pool was generated by ENSEMBLE. ^45^ Finally, SH3-3 originated from a mixed pool containing conformations from both TraDES and ENSEMBLE.

The ELViM projection, containing all the 288 conformations from these three ensembles, is depicted in Figure 4. Although there are much fewer data points in this case, ELViM is capable of generating an effective phase space in which the radius of gyration still varies smoothly throughout the projection. Interestingly, the most unfolded conformations (red dots) form a tail-like structure almost detached from the projection in opposition to the most collapsed ones (dark blue dots).

**Figure 4:**
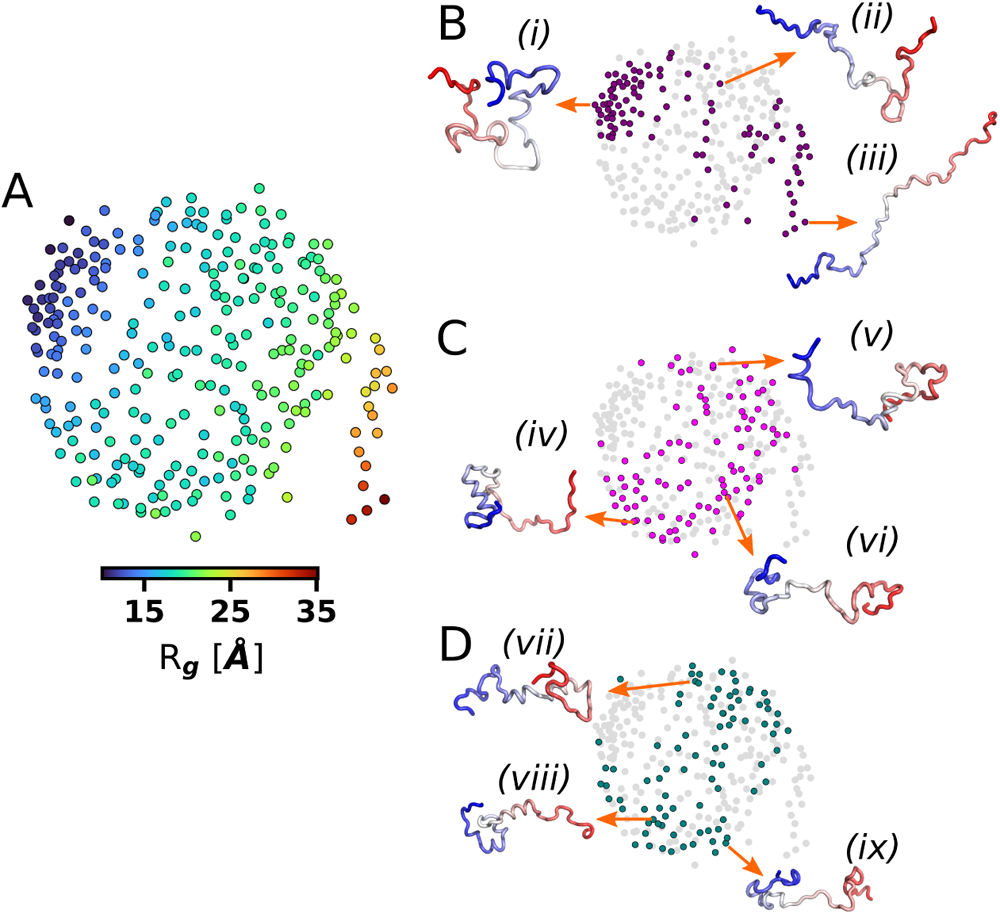
ELViM analysis for the N-terminal SH3 domain of Drk (1-59). (A) ELViM projection generated for the three ensembles of the N-terminal SH3 domains, totaling 288 conformations. Data points are color-coded based on Rg values. The conformations of each ensemble are displayed in the middle column in (B) purple, for SH3-1, (C) fuchsia, for SH32, and (D) teal, for SH3-3. For each case, some arbitrarily chosen structures are displayed in cartoons with the N-terminal in blue.

Contrary to the first analyzed systems, estimating the density of states reliably becomes challenging with few or sparse data points. Nonetheless, a careful analysis of the Figure 4.A reveals some local groups with (almost) superimposed dots. Figures 4.B, C, and D show the individual ensembles for SH3-1, SH3-2, SH3-3, respectively. As an illustration, some conformations were arbitrarily selected and shown in cartoons.

The comparison of the three ensembles shows that SH3-1 (Figure 4B) has a sampling pattern very different from the other ensembles, being formed for most of the more extended and more compact conformations, and only a few conformations from the central portions of the projection. On the other hand, the ensembles depicted in Figures 4C and 4D populates the same region of the ELViM projection, suggesting a similar sampling. However, it is possible to identify differences in the data point density and in the occurrence of small local groups. As for previous cases, a reproducibility analysis is shown in Figure S7 in the Supporting Information.

### Conclusions

Previous studies with ELViM were carried out using strictly molecular dynamics trajectories, and they have proven useful in the investigation of molecular mechanisms, identifying transition-state ensembles, folding routes, and metastable states. The analyzed systems encompass ordered proteins,^39,49,55^ knotted proteins,^51^ disordered peptides and proteins,^50,54,56^ and an RNA tetraloop.^53^ In the present study, we have addressed IDP ensembles that were obtained by a variety of integrative experimental and computational methods. Through the same systematic procedure, this proof of concept has provided valuable insights into their conformational heterogeneity and sampling behavior. The effective phase space generated by ELViM allows for the comparison of different ensembles, highlighting differences in sampling patterns and identifying the most prevalent conformations within each ensemble. This approach facilitates a deeper understanding of the energy landscape of intrinsically disordered protein ensembles, shedding light on their functional mechanisms.

It is difficult to compare ELViM with other analogous approaches mentioned previously, such as those outlined in references.^29,30,32–34^ This difficulty arises due to the myriad of parameters and scores involved in evaluating multidimensional representations.^57,58^ Traditional quality metrics primarily gauge the optimization procedure’s efficiency, falling short in quantifying how effectively the protein conformational space is depicted in the final outcome.^57^ The advantage of using ELViM is the robustness of its representation, but still with adjustable parameters that can deliver an intuitive representation of the energy landscape.

Future work could focus on expanding the application of ELViM to larger data sets of IDP ensembles and exploring its utility in combination with other analysis techniques. For instance, AlphaFold provides only one of the multiple protein conformations and does not provide predictions for intrinsically disordered or unstructured regions. ^59^ ELViM might be integrated into AlphaFold to predict the most likely disordered ensembles. Additionally, the integration of experimental data with ELViM could further enhance the accuracy and reliability of the generated effective phase space. Overall, ELViM proves to be a valuable tool in the study of intrinsically disordered proteins, offering a unique perspective on their conformational landscapes.

## Acknowledgment

We thank Sameer Velankar and Jorge Chahine for their helpful comments and discussions. The authors acknowledge financial support from Prope/Unesp and the Brazilian agencies FAPESP (Grants 2023/02219-1, 2022/08738-8, 2023/08101-2, and 2021/15028-4) and National Council for Scientific and Technological Development – CNPq (Grant 310017/2020-3).

