## Supporting Information for "Understanding the Energy Landscape of Intrinsically Disordered Protein Ensembles"

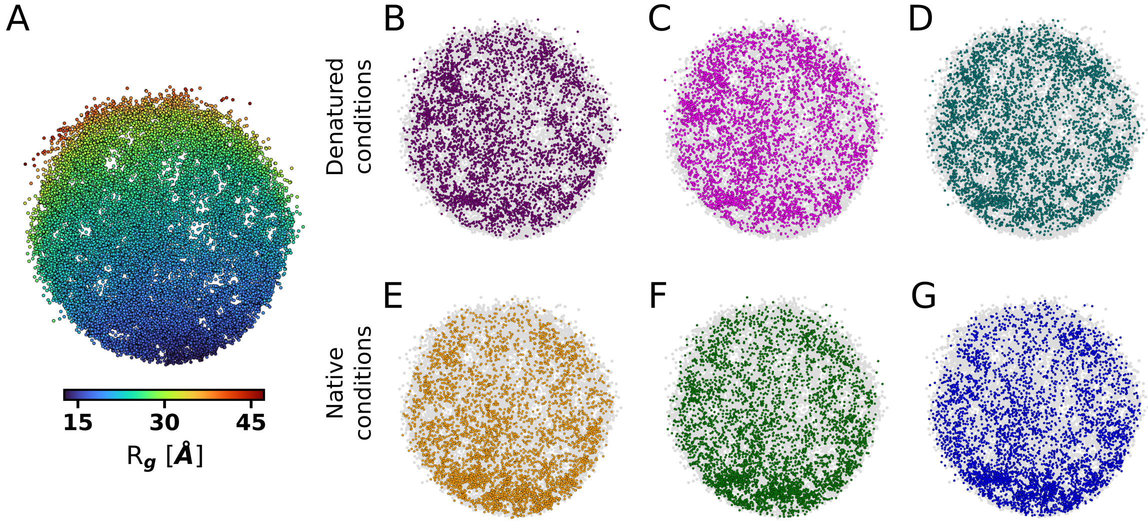


Figure S1: Individual ensembles for NUS. A) ELViM effective phase space color-coded based on the radius of gyration. Dots corresponding to structures from each ensemble — B) NUS1, C) NUS2, D) NUS3, E) NUS4, F) NUS5, G) NUS6 — are depicted in solid color over the entire phase space represented in gray.

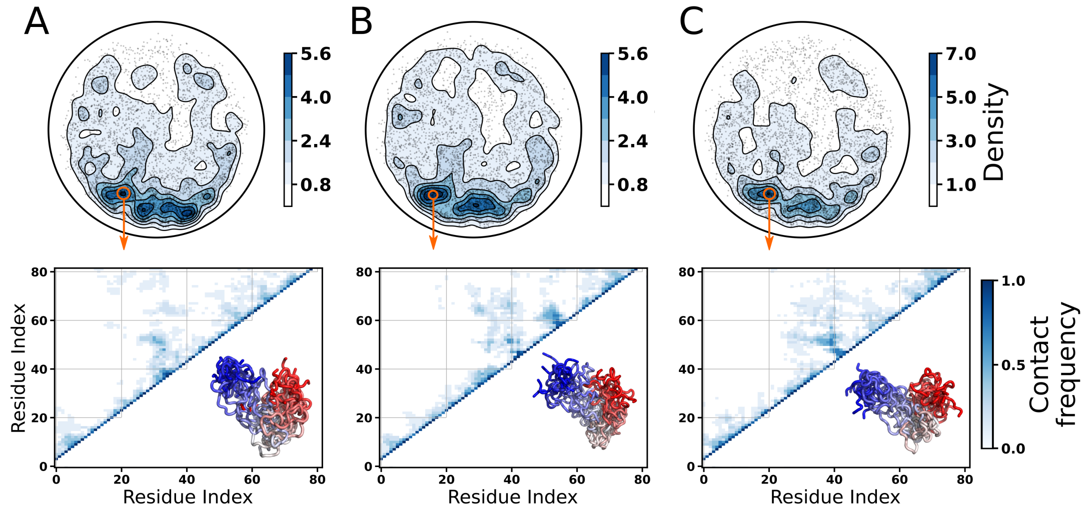


Figure S2: Contact maps for an LCS in the NUS ensembles obtained under native conditions. In order to highlight the consistency, and also the variations, we selected an LCS (10 conformations) from the density peak region encircled in orange. Contact maps, based on a cut-off distance of 10 Å, and the ten superimposed structures are shown in lower panels for (A) NUS4, (NUS5), and (NUS6).

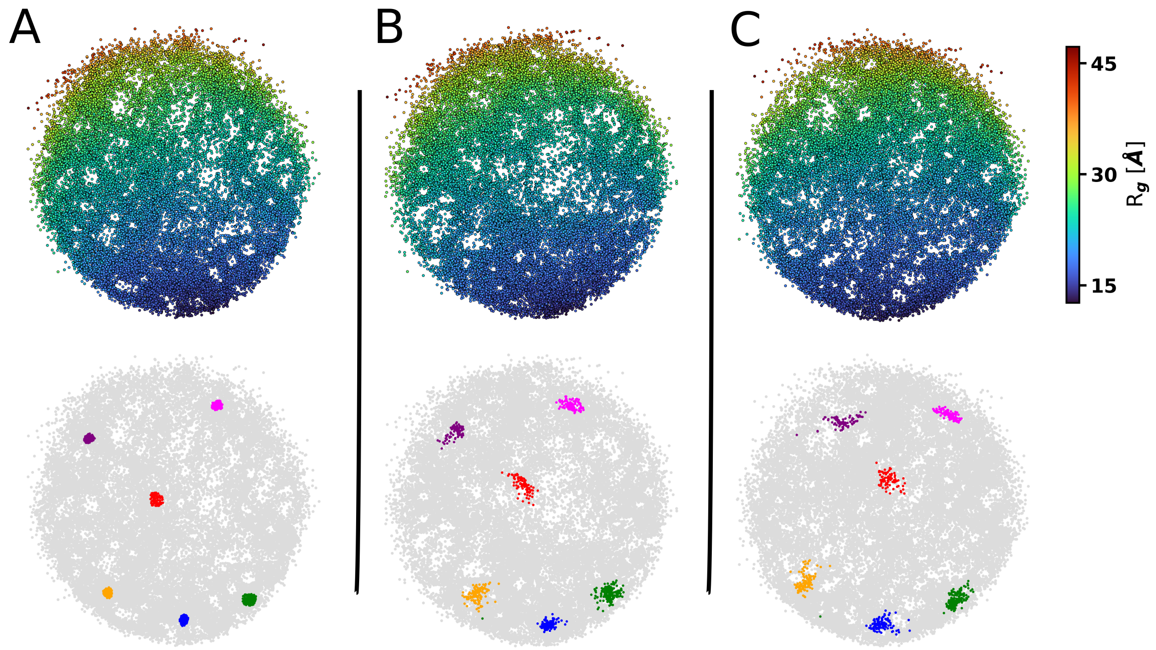


Figure S3: Reproducibility for the NUS ensembles. (A) The same projection shown in the main text is reproduced in the upper panel. Six groups of conformations are arbitrarily selected are depicted in different colors. Two new and independent projection are shown in panels (B) and (C), color-coded based on radius of gyration values (upper panels). In addition, the same groups selected in (A) are shown in lower panel using the same colors. Projections were aligned to facilitate visual comparison.

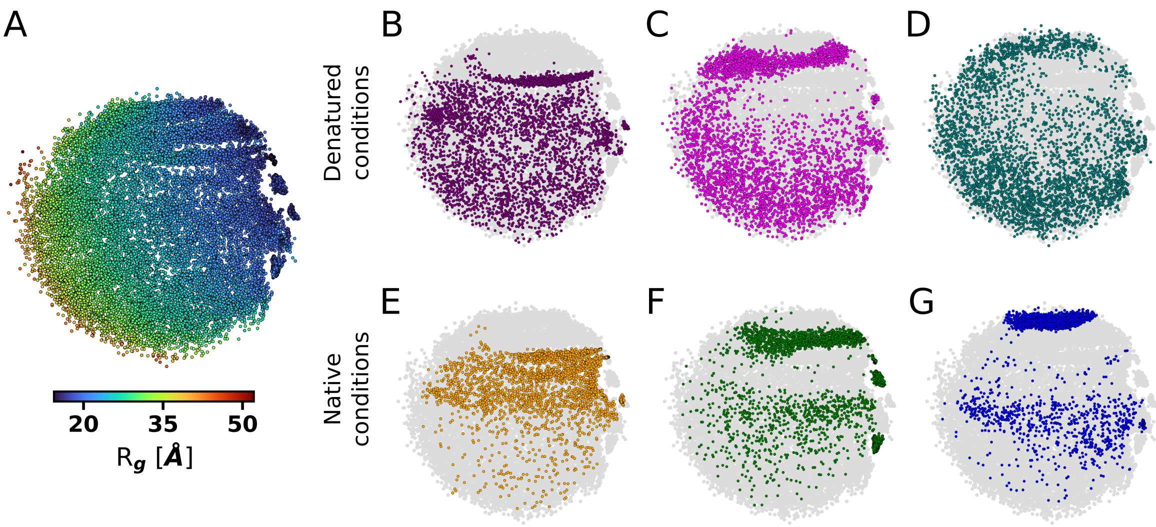


Figure S4: Individual ensembles for NUL. A) ELViM effective phase space color-coded based on the radius of gyration. Dots corresponding to structures from each ensemble — B) NUL1, C) NUL2, D) NUL3, E) NUL4, F) NUL5, G) NUL6 — are depicted in solid color over the entire phase space represented in gray.

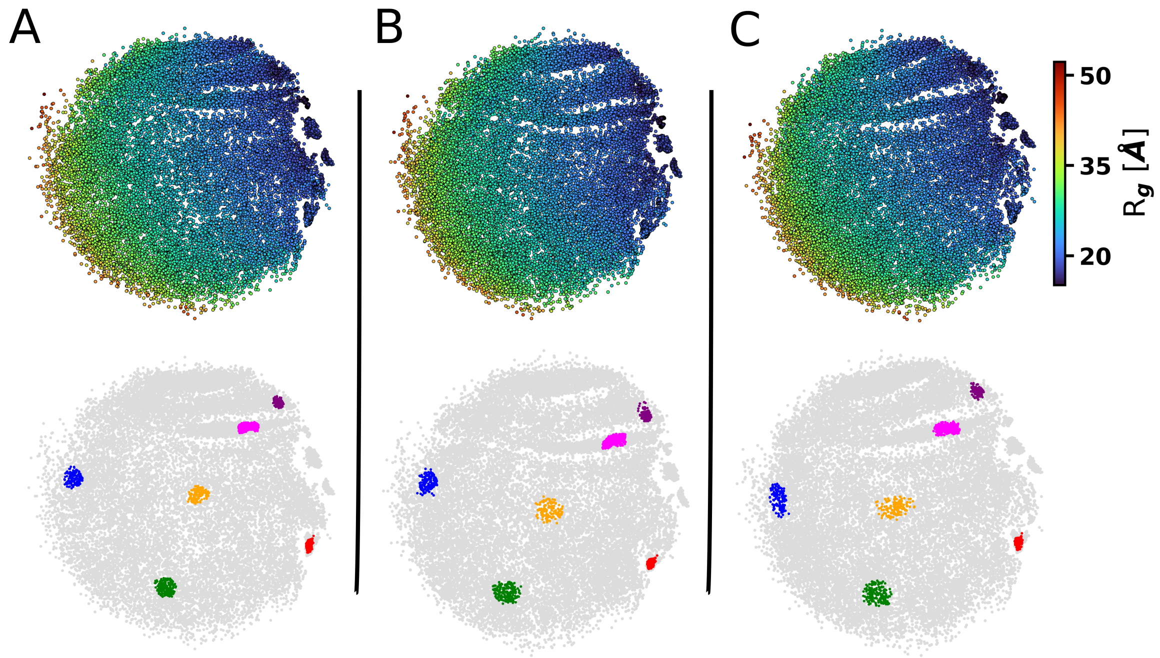


Figure S5: Reproducibility for the NUL ensembles. (A) The same projection shown in the main text is reproduced in the upper panel. Six groups of conformations are arbitrarily selected are depicted in different colors. Two new and independent projection are shown in (B) and (C), color-coded based on radius of gyration values in upper panels. In addition, the same groups selected in (A) are shown in lower panel using the same colors. Projections were aligned to facilitate visual comparison.

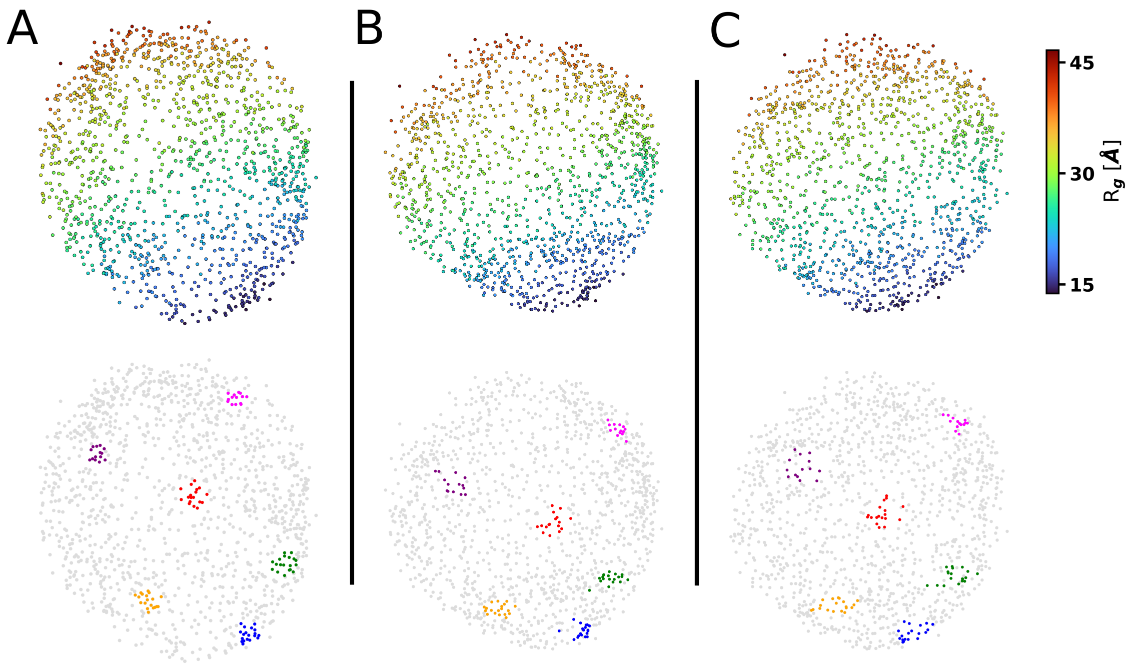


Figure S6: Reproducibility for the Sic1 ensembles. (A) The same projection shown in the main text is reproduced in the upper panel. Six groups of conformations are arbitrarily selected are depicted in different colors. Two new and independent projection are shown in panels (B) and (C), color-coded based on radius of gyration values (upper panels). In addition, the same groups selected in (A) are shown in lower panel using the same colors. Projections were aligned to facilitate visual comparison.

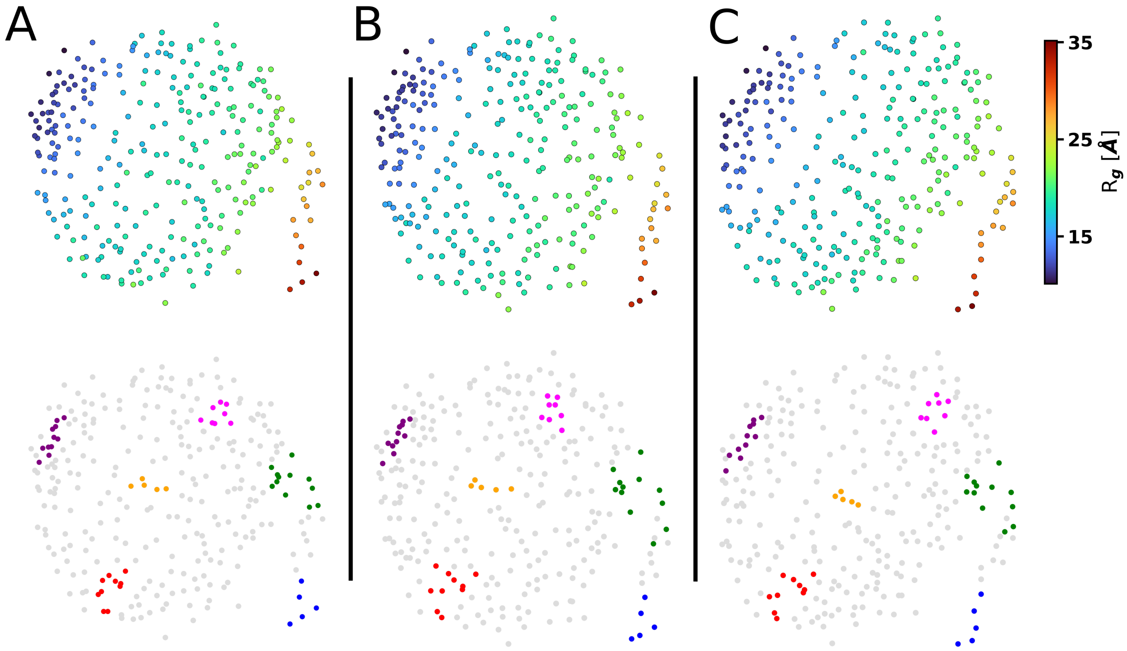


Figure S7: Reproducibility for the N-terminal SH3 domain of Drk ensembles. (A) The same projection shown in the main text is reproduced in the upper panel. Six groups of conformations are arbitrarily selected are depicted in different colors. Two new and independent projection are shown in panels (B) and (C), color-coded based on radius of gyration values (upper panels). In addition, the same groups selected in (A) are shown in lower panel using the same colors. Projections were aligned to facilitate visual comparison.
